# Dynamic pseudo-time warping of complex single-cell trajectories

**DOI:** 10.1101/522672

**Authors:** Van Hoan Do, Mislav Blažević, Pablo Monteagudo, Luka Borozan, Khaled Elbassioni, Sören Laue, Francisca Rojas Ringeling, Domagoj Matijević, Stefan Canzar

## Abstract

Single-cell RNA sequencing enables the construction of trajectories describing the dynamic changes in gene expression underlying biological processes such as cell differentiation and development. The comparison of single-cell trajectories under two distinct conditions can illuminate the differences and similarities between the two and can thus be a powerful tool. Recently developed methods for the comparison of trajectories rely on the concept of dynamic time warping (dtw), which was originally proposed for the comparison of two time series. Consequently, these methods are restricted to simple, linear trajectories. Here, we adopt and theoretically link arboreal matchings to dtw and propose an algorithm to compare complex trajectories that more realistically contain branching points that divert cells into different fates. We implement a suite of exact and heuristic algorithms suitable for the comparison of trajectories of different characteristics in our tool Trajan. Trajan automatically pairs similar biological processes between conditions and aligns them in a globally consistent manner. In an alignment of singlecell trajectories describing human muscle differentiation and myogenic reprogramming, Trajan identifies and aligns the core paths without prior information. From Trajan’s alignment, we are able to reproduce recently reported barriers to reprogramming. In a perturbation experiment, we demonstrate the benefits in terms of robustness and accuracy of our model which compares entire trajectories at once, as opposed to a pairwise application of dtw. Trajan is available at https://github.com/canzarlab/Trajan.

## 1 Introduction

Single-cell RNA sequencing (scRNA-seq) has allowed the detailed dissection of biological processes such as differentiation, development and cell reprogramming. By describing the trajectories along which cells transition to achieve specific cell fates, scRNA-seq can illuminate the dynamic changes in gene expression underlying these processes [19]. Much can be learned from the comparative analysis of single-cell trajectories. Comparing gene expression dynamics along trajectories from two conditions can aid in elucidating the key differences between them and the regulatory programs underpinning the process. For example, comparing the trajectories underlying a given differentiation process in two species would shed light onto the evolutionary differences between these organisms. Comparing the trajectory defining a normal developmental process to that affected by a particular mutation would yield insights into disease mechanisms. Recently, methods have been developed for this purpose, which make use of dynamic time warping (dtw).

Dynamic time warping is a class of algorithms for comparing two time series that advance at different speeds [22]. It was originally developed in the context of automatic speech recognition, but has gained increasing popularity more recently in the comparison of single cell trajectories [1, 6, 9]. Similar to a pairwise sequence alignment that allows for insertions and deletions, dtw finds a mapping (warping) between similar elements in the two sequences to overcome locally stretched and compressed sections. In single-cell trajectories, cells are ordered along pseudo-time and can be aligned based on the expression values of (a subset of) their genes to establish a common pseudotime axis along which expression kinetics become comparable between different conditions.

Dynamic time warping can only compare two time series at a time, and thus current methods for comparing single-cell trajectories are limited to linear trajectories or rely on picking the correct path from a complex trajectory. Even though not based on dtw, MATCHER [23] is similarly restricted to two linear trajectories, built from transcriptomic and epigenetic measurements. It is relevant to mention that complex cell trajectories are common in developmental processes and also arise in response to genetic perturbations [18]. In these cases, prior information such as a set of defined markers would be necessary to pick the most relevant path, but this information is often not available. Another potential caveat of dtw is that it ignores cells that lie on alternative paths and could potentially amplify the signal used to infer the mapping between trajectories.

We present Trajan, a novel method to compare and align complex trajectories with multiple branch points diverting cells into alternative fates (Fig. 1). Trajan automatically identifies the correspondence between biological processes in two trajectories and aligns all of them simultaneously, taking into account their overlap. Given that cells that diverted into different fates share a common ancestry, they cannot be treated as independent from each other. Their independent pairwise alignment (using dtw) could introduce inconsistencies with respect to the mapping of common progenitor cells. Akin to the extension of pairwise alignments to multiple sequence alignment, we seek the best alignment between all corresponding pairs of paths that agree on common progenitor cells. To this end, Trajan adopts arboreal matchings [5] to capture globally consistent similarities between trajectories. Arboreal matchings were originally proposed in the context of phylogentic trees and here we theoretically link them to dynamic time warping. We develop a suite of exact and heuristic algorithms that are suitable for the comparison of trajectories of different characteristics. When aligning single-cell trajectories describing human muscle differentiation and myogenic reprogramming, Trajan automatically identifies the core paths from which we are able to reproduce recently reported barriers to reprogramming. In a perturbation experiment, Trajan correctly maps identical cells in a global view of trajectories, as opposed to a pairwise application of dtw.

**Figure 1:**
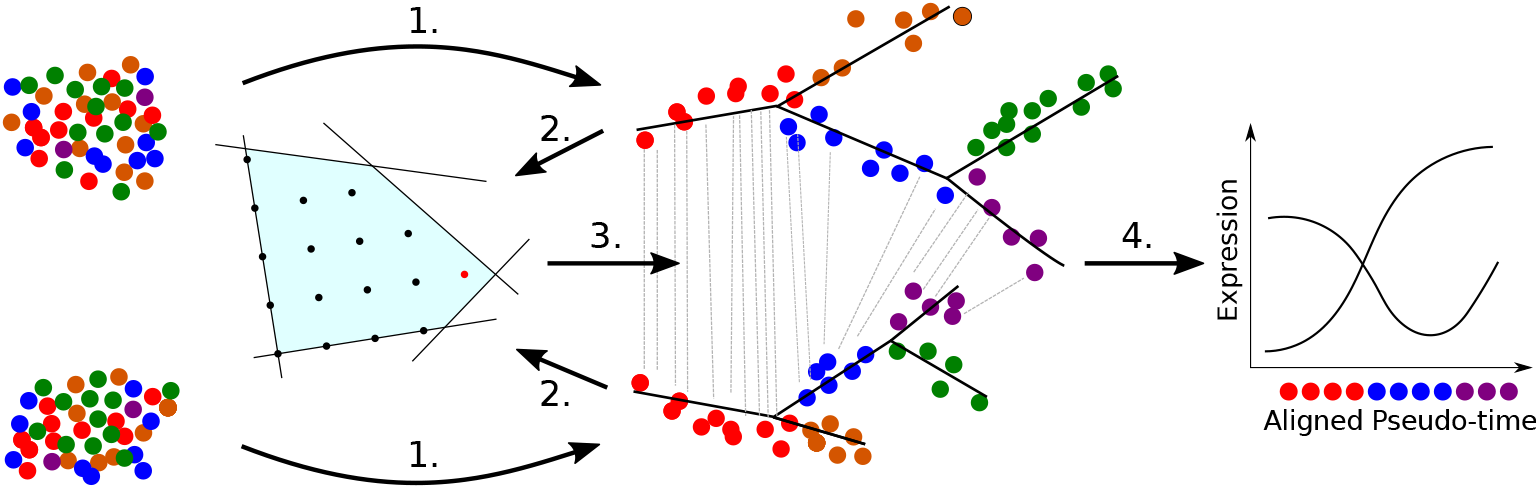
Trajan workflow. 1. Complex trajectories are reconstructed from single-cell RNA measurements using, e.g., Monocle 2. After smoothing and scaling (2.), Trajan aligns entire trajectories by computing an arboreal matching using a branch-and-cut approach (3.), which transforms (warps) the individual pseudo-time scales into a shared one along which expression kinetics can be compared (4.). For simplicity, only the alignment between one pair of paths is shown.

## 2 Methods

Dynamic time warping is the algorithmic workhorse underlying current methods that compare linear single-cell trajectories. In the next section we briefly review the concept of dynamic time warping and show that an attempt to generalize dtw to complex trajectories naturally leads to arboreal matchings between trees, which we have introduced previously in the context of phylogenetic trees [5]. Proofs of Theorems and Lemmas can be found in the Appendix, Section 6.1.

### 2.1 DTW versus arboreal matching

As in classical sequence alignment, dtw matches similar elements in two sequences while preserving their order. To account for different speeds at which the two sequences advance, however, each element of one sequence can be mapped to multiple elements in the other sequence (Fig. 2 left). More formally, given two time series 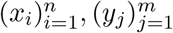, and a distance or similarity measure *d*(*x_i_, y_j_*) ≥ 0 between the time points *x_i_* and *y_j_*, a *warping* is a sequence *p* = (*p*_1_,…,*p_L_*) with *p_ℓ_* = (*n_ℓ_, m_ℓ_*) ∈ [1 : *n*] × [1 : *m*] for *ℓ* ∈ [1 : *L*] that satisfies the following three conditions: (i) *Boundary*: *p*_1_ = (1,1) and *p_L_* = (*n, m*). (ii) *Monotonicity*: *n*_1_ ≤ *n*_2_ ≤ ⋯ ≤ *n_L_* and *m*_1_ ≤ *m*_2_ ≤ ⋯ ≤ *m_L_*. (iii) *Step size*: *p*_*ℓ*+1_ − *p_ℓ_* ∈ {(1, 0), (0,1), (1,1)} for *ℓ* ∈ [1 : *L* − 1]. Note that the example warping in Figure 1 (left) contains no pair of crossing edges and thus preservers the order of the two sequences.

**Figure 2:**
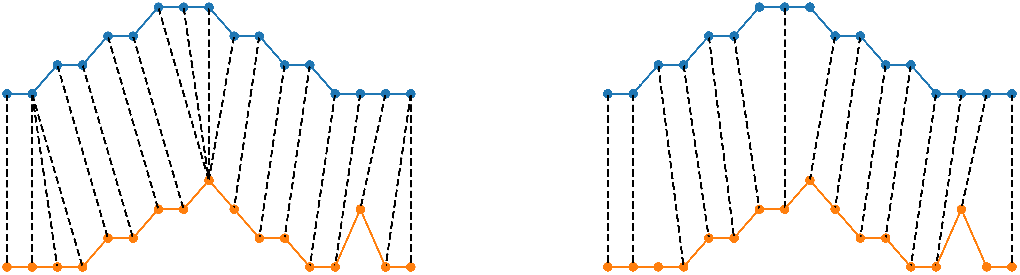
An example of a warping (left) and an arboreal matching (right) between two time series.

The classic dtw aims to find a warping p minimizing the total distance between mapped elements:

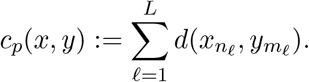

The optimal warping can be computed by a dynamic program that solves:

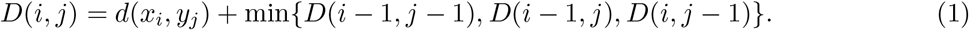

There are various extensions of the classic dtw described above that can be mainly classified as 1) restricting the range of the mapping to a certain window; 2) assigning different weights to different types of steps; and 3) using different step patterns, e.g. *p*_*ℓ*+1_ − *p_ℓ_* ∈ {(1,1), (1,2), (2,1)} in (iii). In the following, we consider the widely used classic dtw which is also the default scheme for computing dtw [24]. Since state-of-the-art methods like Monocle 2 [20] and DPT [11] aim to construct smooth trajectories, the classic dtw provides the necessary flexibility for most single-cell alignment tasks.

Here, we propose a generalization of classic dtw from paths, i.e., linear trajectories, to trees, i.e., complex trajectories: We want to align each path in tree *T*_1_ to at most one path in *T*_2_ and vice versa and, similar to dtw, preserve the order of nodes along the paths, i.e., no crossing edges. In addition, we require all alignments to be consistent, that is, every node must be matched to the same node in all pairwise alignments it is part of. In [5], we have introduced *arboreal matchings* that formalize such a consistent path-by-path alignment of trees: An arboreal matching is a matching *M*, i.e., one-to-one correspondence between nodes in trees *T*_1_ and *T*_2_ such that for any (*u*_1_, *v*_1_), (*u*_2_, *v*_2_) ∈ *M*, *u*_2_ is a descendant of *u*_1_ iff *v*_2_ is a descendant of *v*_1_.

In contrast to dtw, an arboreal matching *M* matches each node (cell) to at most one similar node (cell) in the other tree (trajectory). It is not required to cover all nodes between each pair of paths, but we can flexibly penalize nodes that remain unmatched by *M* in the objective function:

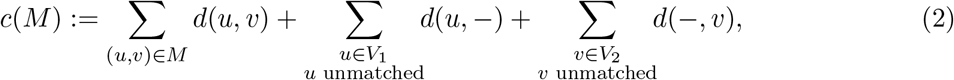

where the cost of leaving node *u*(*v*) unmatched is *d*(*u*, −) > 0 (*d*(−, *v*) > 0). In fact, the arboreal matching of minimum cost (2) between two paths *P* = (*x*_1_,…, *x_n_*) and *Q* = (*y*_1_,…, *y_m_*) can be solved by a very similar dynamic program as in dtw (1):

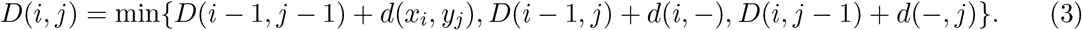

An example arboreal matching between two paths is shown in Figure 2 (right). Again, the noncrossing edges align the two time-series to reveal similarities and unmatched nodes indicate compressed or stretched sections. This makes arboreal matchings as flexible as dtw in the comparison of two trajectories. More specifically, we will show that by choosing an appropriate penalty for unmatched vertices, the optimal dtw and the optimal arboreal matching yield similar measures of similarity or distance of the compared trajectories. Denote by *d_dtw_* and *d_M_* the optimal value of the classic dtw and the arboreal matching between two paths *P* and *Q*, respectively. The following theorem provides an upper bound on *d_dtw_*.

#### Theorem 1.

*Let* 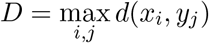. *If* 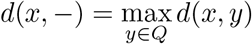 *and* 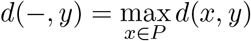, *then*

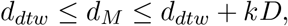

*where k is the minimum number of edges that need to be removed to transform the optimal warping to an arboreal matching*.

Next, we develop a lower bound theorem for the classic dtw. An edge (*x, y*) in the warping *p* is called *redundant* if both vertices *x* and *y* are covered by at least two edges in *p*.

#### Lemma 1.

*There exists an optimal warping of the classic dtw without redundant edges*.

Given an optimal warping *p*\*, we assign penalties to unmatched vertices such that *d_M_* ≤ *d_dtw_*. Let *p*\* be an optimal warping without redundant edges, define

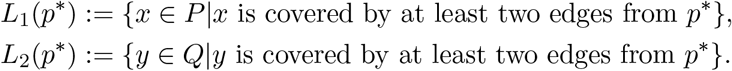

Then, we impose penalties

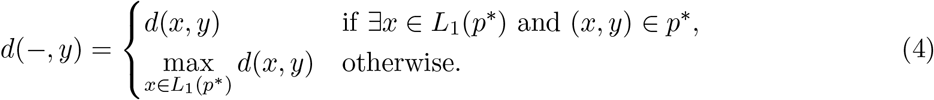

We define penalties *d*(*x*, −), *x* ∈ *P*, analogously. Since *p*\* has no redundant edges, *d*(−, *y*) is uniquely defined. Conversely, if there exist *x*_1_, *x*_2_ ∈ *L*_1_(*p*\*) such that (*x*_1_, *y*) ∈ *p*\*, (*x*_2_, *y*) ∈ *p*\*, the non-redundancy of *p*\* is violated. We have the following lower bound theorem.

#### Theorem 2.

*If d*(*x*, −), *d*(−, *y*) *are defined as in* (4), *then d_M_* ≤ *d_dtw_*.

In Section 3.1 we illustrate how closely the optimal arboreal matchings based on lower and upper bound penalty scheme follow the optimal dtw path.

### 2.2 Limitations of the naïve ILP formulation

Finding the matching minimizing (2) can be phrased as a maximum matching problem that explicitly forbids the two possible types of ancestry violations: Two edges can be crossing, or two nodes on the same root-to-leaf path are matched to nodes on different root-to-leaf paths (Figure 3). The former constraint is equally imposed by dtw, the latter is a consequence of the simultaneous comparison of multiple paths and prevents arbitrary jumps between biological processes in the comparison. In our proof-of-concept study [5], we describe feasible arboreal matchings between two rooted trees *T*_1_ = (*V, E*_1_), *T*_2_ = (*V*_2_, *E*_2_), by the following simple ILP:

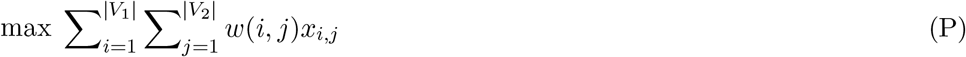

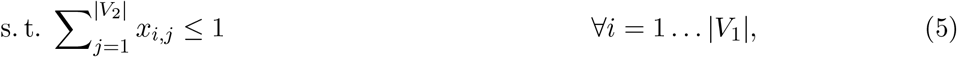

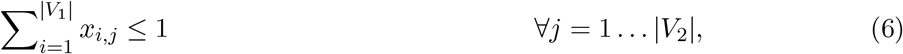

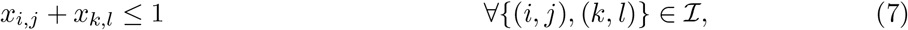

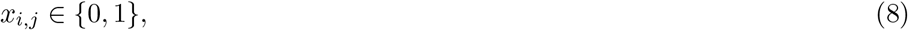

where indicator variables *x_i,j_* denote the presence or absence of an edge (*i,j*), weights *w*(*i,j*) := *d*(*i*, −) + *d*(−, *j*) − *d*(*i, j*). Pairs of edges (*i, j*) and (*k, l*) are *compatible* if it holds that *k* is a descendant of *i* in *T*_1_ iff *l* is a descendant of *j* in *T*_2_. Set 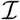 contains pairs of edges {(*i, j*), (*k, l*)} that are incompatible, i.e., they are either crossing or one-sided independent (Figure 3).

**Figure 3:**
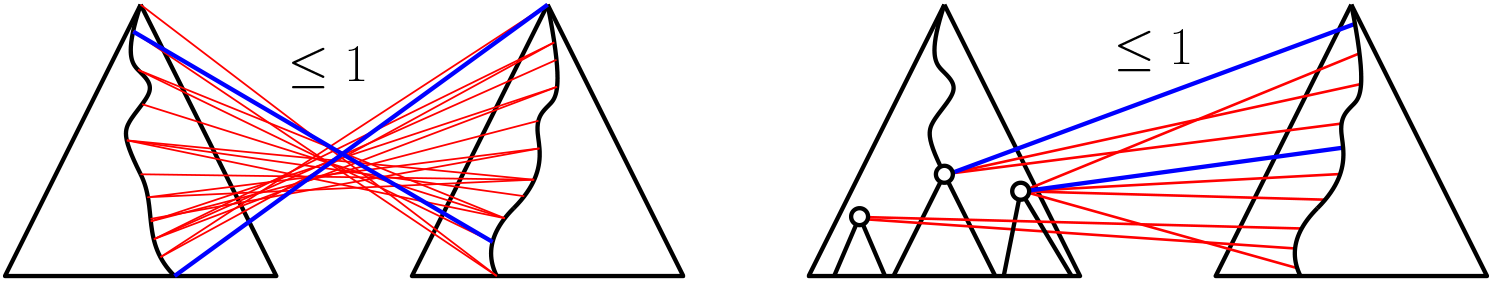
Pair of crossing edges (blue) extended to a clique of crossing edges (left) and pair of semi-independent edges (blue) extended to a clique of semi-independent edges (right).

As our experiments in Section 3.3 show, this ILP formulation does not allow to practically align trajectories comprising as few as 100 single cells. In the following theorem, we identify its weak LP-relaxation as a theoretical explanation for this empirical performance, since the search space that needs to be explicitly explored by an ILP solver depends on the strength of the LP relaxation. Let OPT denote an optimal solution to the above ILP and let *w*(OPT) be its optimal score. Let |*V*_1_| = *n*, |*V*_2_| = *m*, and w.l.o.g we assume *n* ≤ *m*.

#### Theorem 3.

*The integrality gap of the linear programming relaxation of* (P) *is n* − *o*(1).

### 2.3 A branch-and-cut algorithm for arboreal matchings

In this section, we introduce a thoroughly engineered branch-and-cut algorithm that allows to practically compare complex single-cell trajectories. Its main ingredients are (i) cuts that trim the LP relaxation closer to the convex hull of feasible arboreal matchings (Section 2.3.1), (ii) polynomial-time algorithms that can find these cuts on demand (Section 2.3.2), (iii) a branch-and-bound scheme that makes use of modern CPU architectures (Section 2.3.3), and (iv) an in-house developed, non-commercial, non-linear solver that we use for all continuous optimization problems (Appendix, Section 6.3). In Section 6.5 in the appendix, we generalize arboreal matchings and our branch-and-cut algorithm to directed acyclic graphs.

#### 2.3.1 Valid clique constraints

The next theorem motivates the addition of valid inequalities to reduce the integrality gap of the LP relaxation. Let (PC) be the LP-relaxation of the ILP that we obtain by replacing the pairwise incompatibilities (7) in (P) by the more general clique inequalities

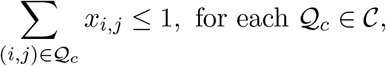

where 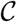 is a set of cliques, i.e., sets of pairwise incompatible edges. Denote by *E*(*T*_1_, *T*_2_) the set of edges between two trees. For each *M* ⊂ *E*(*T*_1_, *T*_2_), let *r*(*M*) denote the maximum size of any feasible (unweighted) arboreal matching contained in *M*.

##### Theorem 4.

*If we can write any set M* ⊂ *E*(*T*_1_, *T*_2_) *as union of at most r*(*M*) *cliques, that is*, 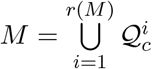, *then the integrality gap of the linear program* (PC) *is at most* log *n* + 1.

Inspired by this theorem, we strengthen the LP relaxation by lifting pairwise incompatibility constraints (7) to maximal sets (cliques) of pairwise crossing edges and maximal sets (cliques) of edges that are pairwise semi-independent (Figure 3). As our experiments in Section 3.3 indicate, these lifted constraints result in stronger bounds that allow us to prune larger parts of the search space. Due to their exponential number, we add lifted constraints only on demand, that is, if they cut off the current optimal fractional solution. In the next section, we describe how to assess this demand in polynomial time.

#### 2.3.2 Polynomial-time separation algorithms

In the following, we consider trees *T*_1_ = (*V*_1_, *E*_1_) and *T*_2_ = (*V*_2_, *E*_2_) with roots *r*_1_ and *r*_2_, sets of leaves 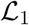 and 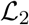, and parent mappings *π*_1_ and *π*_2_, respectively. For two vertices *p, q* ∈ *V_i_*, denote by [*p, q*] the unique path in *T_i_* between *p* and *q*.

##### Crossing edges clique constraints

A maximal set of pairwise crossing edges between two fixed root-to-leaf paths [*r*_1_, *ℓ*_1_], [*r*_2_, *ℓ*_2_] can be obtained by the following procedure. Starting from an edge between leaf *ℓ*_1_ in *T*_1_ and the root of *T*_2_, in each step we either move up along [*r*_1_, *ℓ*_1_] and keep the node in *T*_2_ fixed, or we move down along [*r*_2_, *ℓ*_2_] and keep the node in *T*_1_ fixed. Analogously we can start from edge (*r*_1_, *ℓ*_2_). Figure 3 (left) shows one possible outcome of this procedure.

Given a fractional solution *x*\* to the current LP relaxation, the separation problem asks to find a hyperplane that cuts off (separates) *x*\* from the polytope without losing any feasible integral solution, i.e., arboreal matching. Since each maximal set (clique) 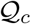 of edges obtained by the procedure above are pairwise incompatible, the sum of their fractional values must not exceed 1:

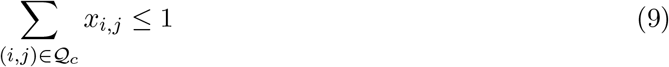

Among the exponentially many crossing cliques 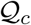 we can identify one for which (9) is (most) violated efficiently by a dynamic program. For fixed paths *P*_1_ = [*r*_1_, *ℓ*_1_], *P*_2_ = [*r*_2_, *ℓ*_2_], let *D*[*u, v*] denote the maximum (with respect to *x*\*) clique between [*r*_1_, *u*] and [*v, ℓ*_2_]. It can be defined recursively as the better choice between moving up in *P*_1_ or down in *P*_2_:

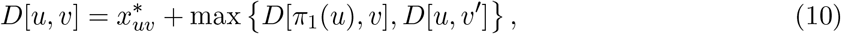

where *π*_2_(*v*′) = *v*. The maximum *x*\*-weight clique *D*[*ℓ*_1_, *r*_2_] can then be computed by a dynamic program in time 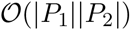. We can generalize this dynamic program to two trees by considering all child nodes *v*′ in the recursion instead of the unique descendant along the path:

#### Theorem 5.

*Given a fractional solution x*\* *we can determine whether a crossing edge clique inequality* (9) *is violated in time* 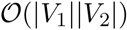.

##### Semi-independent clique constraints

We strengthen the LP relaxation even further by lifting pairs of semi-independent edges to maximal sets (cliques) of pairwise semi-independent edges. Such a set consists of edges that are all incident to nodes on a common root-to-leaf path in one tree, and are incident to nodes in the second tree that all lie on distinct root-to-leaf paths, i.e., are independent (Figure 3 right). Again, edges in such a clique 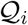 must satisfy

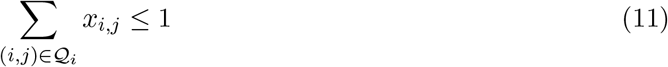

Formally, the separation of semi-independent clique constraints with respect to *T*_1_ and *T*_2_ is given by the following theorem.

###### Theorem 6.

*Given a fractional solution x*\* *we can determine whether a semi-independent clique inequality* (11) *is violated in time* 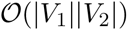.

Again, Theorem 6 allows us to cut off large parts of the polytope in polynomial time, without losing any feasible arboreal matching.

#### 2.3.3 Obtaining integral solutions

In the previous section we described a tighter relaxation of the original ILP formulation. While this relaxed LP improves upon previous approaches it still provides only fractional solutions in the worst case. Trajan implements four different strategies to obtain integral solutions, ranging from fast, but suboptimal, to more expensive, but optimal methods. With this we try to address the need for tailored trade-offs between accuracy and speed imposed by different single-cell sequencing technologies that assay a variable number of genes at varying resolution in hundreds to millions of cells. We describe heuristic approaches implemented in Trajan in Appendix Section 6.2.

##### Branch and bound

Trajan can compute an optimal arboreal matching by a classical branch and bound algorithm, whose running time can be exponential in the worst case. We have implemented several node selection strategies [7], including best first, depth first and a hybrid approach as well as various common variable selection schemes [3], including most and least fractional variables, and most constrained variables. The initial primal bound is obtained by the simple greedy approach described above or the fixed-parameter tractable algorithm described in Section 2.4. Taking advantage of modern CPU architectures, Trajan can run multiple instances of our solver in parallel, each one using a different node and variable selection strategy while sharing the current best primal bound in memory. In best first mode, Trajan can distribute open subproblems across a user-specified number of processors. Note that a tighter relaxation can speed up the running time of the branch and bound exploration since better dual bounds allow for a better pruning of some of the subtrees in the branch and bound computation graph. This can be seen in the experiments in Section 3.3.

### 2.4 FPT algorithm for small number of cell fates

For trajectories with a small number of cell fates *k* we employ a fixed-parameter tractable algorithm, parameterized by *k*. It guesses the correspondence between paths in the two trajectories and applies a dynamic program similar to [13] to align them optimally, in total time 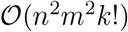, where *k* is the smaller number of leaves among the two trees comprising *n* and *m* nodes, respectively. Even for large *k*, our branch and bound solver optionally runs the FPT algorithm for a small number of path permutations to derive a primal bound on the optimal solution.

## 3 Results

We have implemented the branch-and-cut algorithm described above and have bundled it with our non-linear solver (Section 6.3) in our novel trajectory alignment tool Trajan. In addition to a full branch and bound scheme, Trajan offers heuristic approaches to transform strong bounds into feasible arboreal matchings as well as an FPT algorithm for small number of cell fates at a dramatically reduced computational cost. Trajan adopts a strategy similar to [6] to prepare the output of Monocle 2 (or similar trajectory reconstruction methods) for a meaningful alignment, including the smoothing and scaling of expression curves.

### 3.1 Lower and upper bounds on dtw

Here, we illustrate the practical relevance of the upper bound (UB, Theorem 1) and lower bound (LB, Theorem 2) that the optimal arboreal matching between two paths can provide on the optimal dtw. We align two simple trajectories constructed from scRNA-seq data on dendritic cells stimulated under two conditions (LPS and PAM) collected at 4 time points after stimulation [21]. The two linear trajectories and the dissimilarity matrix were obtained from a recent study that introduced cellAlign [1], a method that aligns two simple trajectories based on dtw. The optimal solutions computed by cellAlign using dtw and by Trajan using the LB penalty scheme are equivalent (Figure 4). When using the UB penalty scheme, Trajan’s optimal path through the dissimilarity matrix roughly follows the dtw path and represents a solution with almost 2 times larger score.

**Figure 4:**
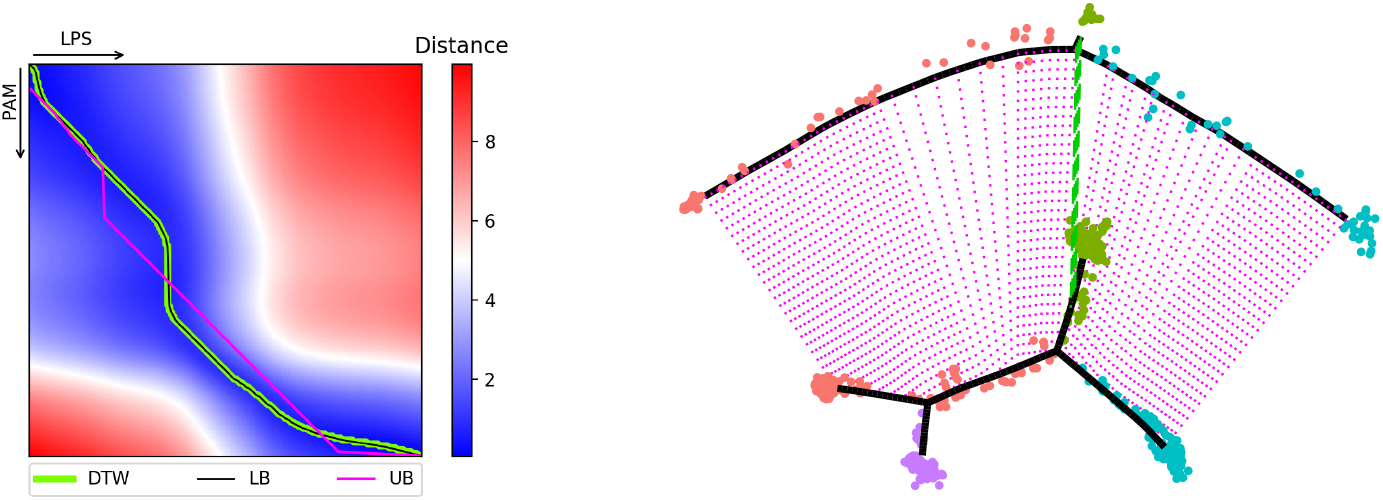
The optimal dtw path and two optimal paths computed by Trajan with lower bound (LB) and upper bound (UB) penalty scheme. The optimal dtw path and Trajan’s LB path coincide (left). Alignment of myogenic reprogramming and differentiation dynamics (right). Trajan discovers the core branches of similar cell fates.

### 3.2 Trajan reproduces barriers in myogenic reprogramming

Here, we re-analyzed two public single-cell datasets: human skeletal muscle myoblast (HSMM) differentiation and human fibroblasts undergoing MYOD-mediated myogenic reprogramming (hFib-MyoD). These datasets were previously analyzed in [6], where the authors set out to compare these related processes in order to identify molecular barriers that hinder the efficient reprogramming of fibroblasts to myotubes. The authors used known myoblast differentiation markers (CDK1, ENO3, MYOG) to identify the core path within the complex trajectory constructed from hFib-MyoD, and they aligned this path to the core path in normal muscle development (HSMM) using dtw. The authors pointed out that the combined trajectory constructed from cells in both conditions did not intermix cells and thus did not allow to assess critical commonalities and differences in expression dynamics. We repeated the single-cell data analysis described in [6] to obtain the corresponding trajectories from Monocle 2 (Figure 5).

**Figure 5:**
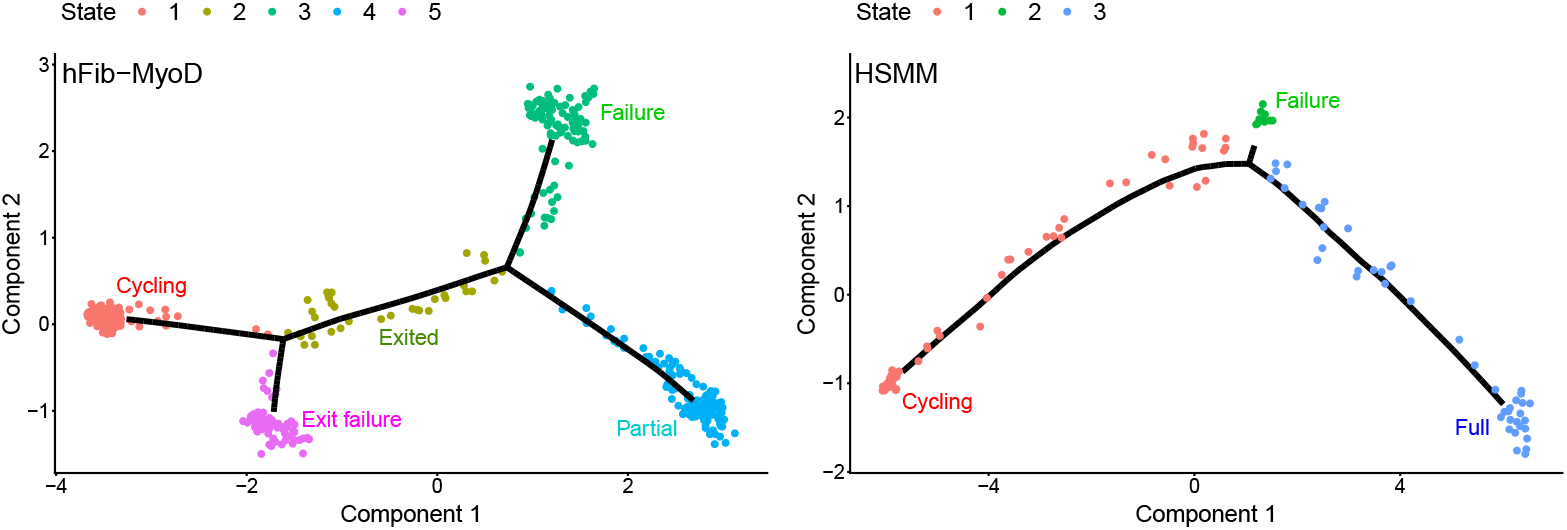
Trajectories of myogenic reprogramming (left) and differentiation (right). Cycling: undifferentiated, actively proliferating cells; Exited: cells lacking expression of cell cycle and muscle contraction genes; Exit failure: cells expressing genes of early myoblast differentiation yet still proliferating; Failure: cells lacking expression of cell cycle genes as well as of muscle contraction genes; Partial: cells expressing MYOG and multiple muscle contraction genes and lacking expression of cell cycle genes; Full: full progression to contractile myotubes.

We then sought to align these complex trajectories using our algorithm. We show that Trajan is able to align the core paths of each complex trajectory, without any previous knowledge or path picking, using the same distance measure (correlation) as in the original publication. The global dynamics alignment of HSMM and hFib-MyoD are shown in Figure 4. Interestingly, our approach not only aligns the core trajectories, but it also aligns the branches corresponding to failure of reprogramming, which are characterized in both processes by cells that exited the cell cycle, yet failed to proceed toward differentiation [6], [19].

After performing the trajectory alignment with Trajan, we constructed gene expression kinetics plots for a set of genes that were assessed in [6] to investigate whether our alignment was able to reproduce their reported findings regarding similarities and differences between these two processes. Indeed, we were able to reproduce their key findings: Proliferation marker CDK1 is downregulated both in HSMM and hFib-MyoD; Muscle transcriptional regulators (MEF2C, MYOG) are upregulated later and to a lesser extent in hFib-MyoD compared to HSMM; BMP4 is only expressed in hFib-MyoD and ID family proteins (ID1, ID3) which lie downstream of BMP signaling fail to be downregulated in hFib-MyoD; IGF pathway genes (IGF2, IGF1R) are expressed at higher levels in HSMM (Figure 6).

**Figure 6:**
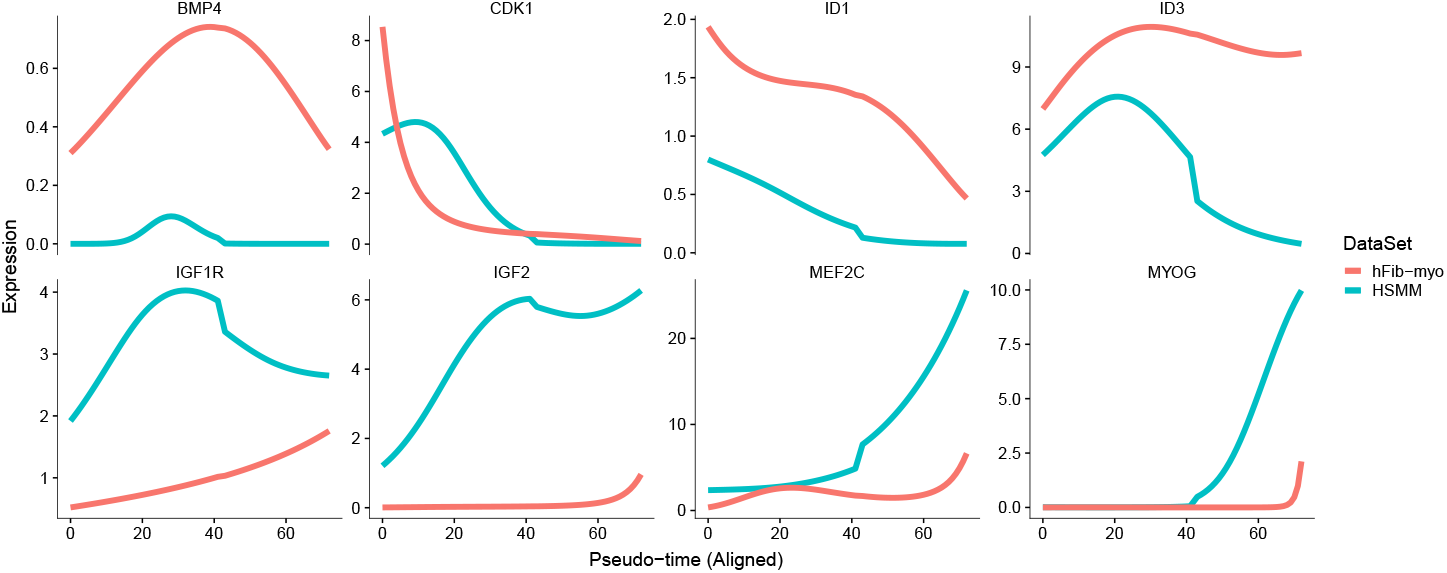
Gene expression dynamics after trajectory alignment with Trajan.

We evaluated Trajan using penalty schemes that assign the maximum and average weight of incident edges as well as the minimum cost implied by the lower bound Theorem 2 over all pairs of paths (*lb*). While the maximum scheme (*max*) is a direct generalization of the cost scheme applied by Theorem 1, the averaging scheme (*avg*) tries to capture the expected cost of leaving a vertex unmatched and is the default scheme applied by Trajan. All schemes correctly picked the correct core paths in the two trajectories and are robust under subsampling. (Appendix, Figure 8).

### 3.3 Accuracy of Trajan

Here, we compare the accuracy of Trajan in matching “correct” cells between complex trajectories to the path-wise alignment by dtw. To this end, we perturb the hFib-MyoD trajectory output by Monocle 2 by randomly subsampling 80% of the input cells and 80% of the genes used for ordering them along pseudo-time. We align isomorphic trajectories (trees) comprising a variable number of nodes (parameter *ncenter* in Monocle 2), measuring the difference between nodes by Euclidean distance. Since we know the true correspondence of nodes between different perturbed trees, we can count false positive and false negative alignments as a measure of accuracy. In Table 1 we report the number of false positive (FP) and false negative (FN) alignments of the classic dtw run on each true pair of paths, and Trajan using different penalty schemes (avg, max, lb). Trajan takes the entire trees as input, it is not given the correct path-to-path correspondence. Nevertheless, Trajan almost always finds the true correspondence between cells, compared to the path-wise dtw scheme, that introduced both FP and FN alignments.

**Table 1:** Average number of false positive (+) and false negative(−) alignments of Trajan and path-wise dtw. The average is taken over a variable number of instances comprising a total # of nodes in the two input trees.

| # of nodes | # of instances | Trajan |  |  |  |  |  | DTW |  |
| --- | --- | --- | --- | --- | --- | --- | --- | --- | --- |
|  |  | avg<sup>+</sup> | avg<sup>-</sup> | max<sup>+</sup> | max<sup>-</sup> | lb<sup>+</sup> | lb<sup>-</sup> | FP | FN |
| 80 | 435 | 0.0 | 0.2 | 0.0 | 0.0 | 12.8 | 17.7 | 35.0 | 32.1 |
| 100 | 435 | 0.0 | 0.0 | 0.0 | 0.1 | 12.9 | 17.6 | 54.6 | 50.5 |
| 140 | 190 | 0.0 | 0.0 | 0.0 | 0.0 | 18.6 | 30.0 | 54.5 | 50.7 |
| 180 | 190 | 0.0 | 0.0 | 0.0 | 0.2 | 31.8 | 38.9 | 83.9 | 76.0 |
| 210 | 45 | 0.0 | 0.0 | 0.0 | 0.0 | 33.9 | 43.9 | 76.6 | 70.3 |

Table 2 reports the running times of the naïve ILP using the commercial solver IBM ILOG CPLEX 12.7 and Trajan coupled with our in-house non-linear solver on a random subset of the instances introduced above. In addition to the full branch-and-cut implementation (Trajan-BnC), we ran Trajan switching to the FPT algorithm (Trajan-FPT). On a 2.30GHz Linux system using up to 15 threads, Trajan-BnC is at least 13 times faster than the naïve ILP using CPLEX, while Trajan-FPT is another 10 times faster then Trajan-BnC. The speedup of Trajan-FPT on this set of instances is not surprising, since these trees comprise only 3 different leaves (cell fates). CPLEX was not able to solve instances with more than 200 nodes since it exceeded the memory limit of 320 GB.

**Table 2:**
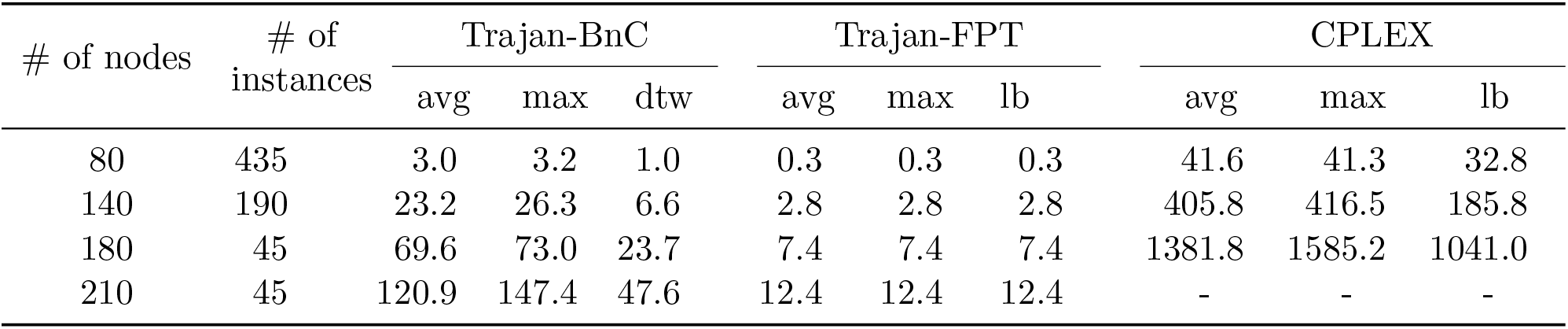
Average runtime in seconds of Trajan vs CPLEX

| # of nodes | # of instances | Trajan-BnC |  |  | Trajan-FPT |  |  | CPLEX |  |  |
| --- | --- | --- | --- | --- | --- | --- | --- | --- | --- | --- |
|  |  | avg | max | dtw | avg | max | lb | avg | max | lb |
| 80 | 435 | 3.0 | 3.2 | 1.0 | 0.3 | 0.3 | 0.3 | 41.6 | 41.3 | 32.8 |
| 140 | 190 | 23.2 | 26.3 | 6.6 | 2.8 | 2.8 | 2.8 | 405.8 | 416.5 | 185.8 |
| 180 | 45 | 69.6 | 73.0 | 23.7 | 7.4 | 7.4 | 7.4 | 1381.8 | 1585.2 | 1041.0 |
| 210 | 45 | 120.9 | 147.4 | 47.6 | 12.4 | 12.4 | 12.4 | - | - | - |

## 4 Conclusion

We have introduced Trajan, a novel method that allows for the first time the alignment of complex (non-linear) single-cell trajectories. Originally introduced to compare phylogenetic trees, in Trajan we adopt arboreal matchings to perform an unbiased alignment enabling the meaningful comparison of gene expression dynamics along a common pseudo-time scale. Trajan does not make any assumptions concerning the algorithm used to reconstruct the trajectory and can in principle be coupled with any available reconstruction method. In a future algorithm, an arboreal matching between cells might prove useful in guiding a joint learning of trajectories for two biological processes. Furthermore, our generalization to directed acyclic graphs (see Appendix) can be used to align data-driven ontologies and the manually curated Gene Ontology (GO) to assign genes to existing GO terms, but also to infer new terms and potentially confirm or correct hierarchical term-term relationships [8].

## 5 Acknowledgments

Sören Laue has been funded by Deutsche Forschungsgemeinschaft (DFG) under grant LA 2971/1-1. Mislav Blaževič was supported in part by BAYHOST. Francisca Rojas Ringeling was supported by the Bavarian Gender Equality Grant (BGF).

## 6 Appendix

### 6.1 Proofs of Theorems and Lemmas

#### Proof of Theorem 1.

(sketch) The first inequality is proven by induction on *i* + *j*.

Let *p*\* be the optimal warping and k the minimum number of edges that need to be removed to transform *p*\* to a feasible arboreal matching *M*. Since *M* has *k* unmatched vertices, we have

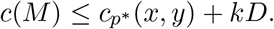

This implies that *d_M_* ≤ *d_dtw_* + *kD*, which also completes the proof of the theorem.

#### Proof of Lemma 1.

Conversely, let *p*\* be an optimal warping such that (*x_i_, y_j_*) ∈ *p*\* and both *x_i_* and *y_j_* are covered by at least two edges in *p*\*. From the coverage property of the warping we must have (*x_i_, y*_*j*−1_) ∈ *p*\* or (*x_i_, y*_*j*+1_) ∈ *p*\*. If (*x_i_, y*_*j*−1_) – *p*\*, we get (*x*_*i*+1_, *y_j_*) ∈ *p*\* since *y_j_* is covered by at least two edges in *p*\* and by the warping conditions. As a results, *D*(*i, j*) = *D*(*i, j* − 1) + *d*(*x_i_, y*_j_) and *D*(*i* + 1, *j*) = *D*(*i, j*) + *d*(*x*_*i*+1_, *y_j_*) = *D*(*i, j* − 1) + *d*(*x_i_, y_j_*) + *d*(*x*_*i*+1_, *y_j_*). From (1), we obtain *D*(*i* + 1, *j*) ≤ *D*(*i, j* − 1) + *d*(*x*_*i*+1_, *y_j_*). This implies that *d*(*x_i_, y_j_*) ≤ 0. Since *d*(*x_i_, y_i_*) > 0 we must have *d*(*x_i_, y_j_*) =0. As a consequence, we can remove (*x_i_, y_j_*) from *p*\* without violating the warping conditions. The case (*x_i_, y*_*j*+1_) ∈ *p*\* is proven in an analogous manner.

#### Proof of Theorem 2.

Let *p*\* be a non-redundant optimal warping. For every vertex *x* ∈ *L*_1_(*p*\*) and *y* ∈ *L*_2_(*p*\*) we delete all incident edges but one, which results in an arboreal matching of the same cost as dtw *p*\*. Hence, it implies that *d_M_* ≤ *d_dtw_*.

#### Proof of Theorem 3.

Let 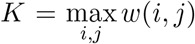, hence *K* is bounded above by *w*(OPT). Moreover, for any feasible solution *x* to the relaxation, we have

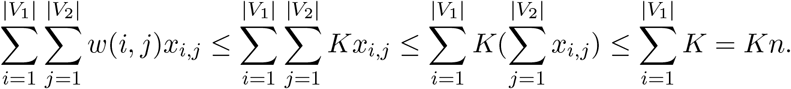

Therefore, the optimal value of the LP relaxation is at most *n* times *w*(OPT). Our bad instance consists of the two rooted trees shown in Figure 7, with *w*(red/blue edges) = 1 and *w*(·) = 0 otherwise. Any pair of nonzero weight edges are incompatible, so the maximum cost matching is 1. Let 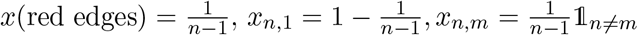, and *x*(·) = 0 otherwise, where 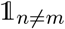 is a binary number such that 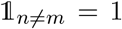 iff *n* = *m*. Hence, *x* is a feasible solution with cost of (*n* − 1)^2^/(*n* − 1) + 1 = *n* if *n* ≠ *m* and 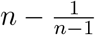 if *n* = *m*. Therefore, the optimal value of the LP relaxation at least *n* − *o*(1).

**Figure 7:**
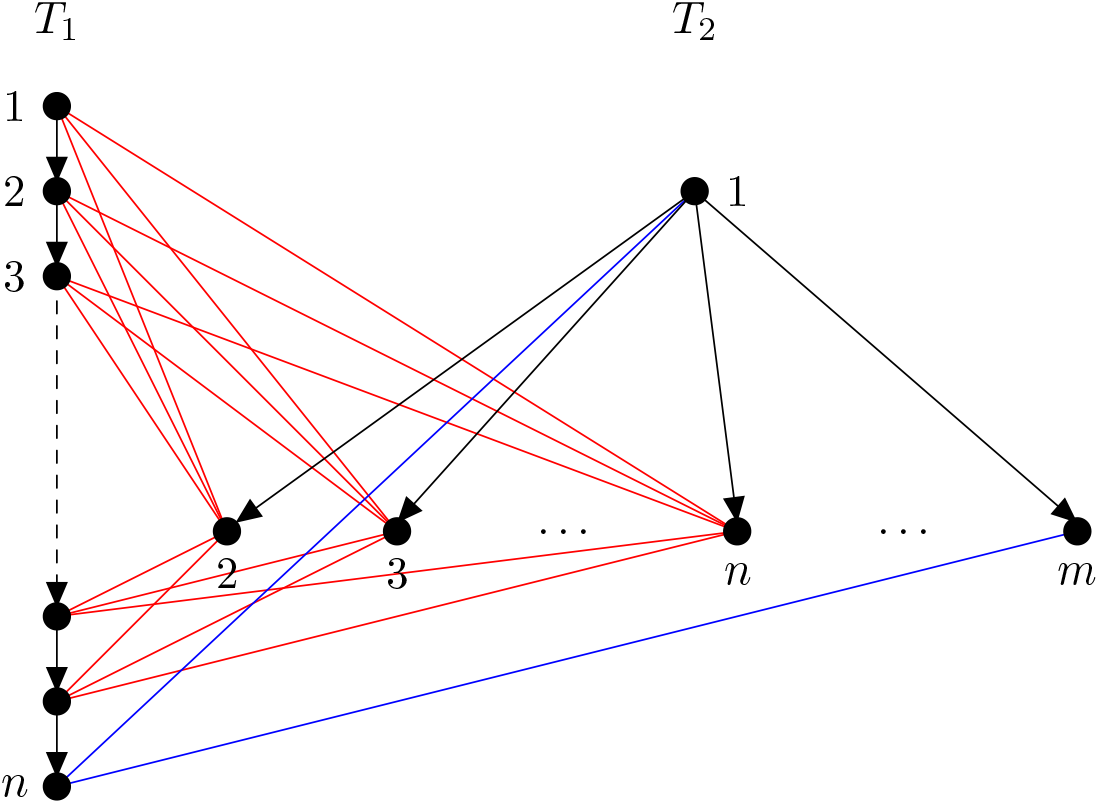
The integrality gap of the LP relaxation of (P) is *n*.

#### Proof of Theorem 4.

Let *x*\* be an optimal solution of the LP relaxation with clique constraints. We show that there exists a feasible arboreal matching with cost at least 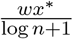, where 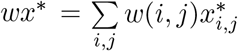. Let *l* be the smallest integer such that for all *e* ∈ *E*(*T*_1_, *T*_2_), 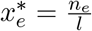, where *n_e_* ∈ ℕ. We will decompose *lx*\* = *x*^1^ + *x^N^*, where *x^i^* is an incident vector (with ground set E) of some feasible arboreal matching. Next, we are going to show that *N* ≤ *l*(log *n* + 1). The procedure for finding *x^i^* is inductive. Let *y^i^* = *lx*\* − (*x*^1^ + ⋯ + *x^i^*) for *i* = 1,2,… and *y*^0^ = *lx*\*. Define

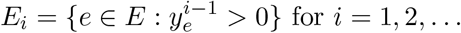

Let *x_i_* be an incident vector of a maximum feasible arboreal matching in *E_i_* such that *x^i^*(*E_i_*) = *r*(*E_i_*). By assumption, we can write 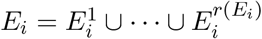, where 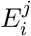 is a clique. As the result, we have

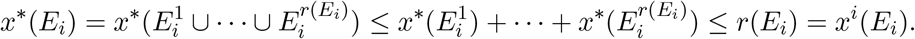

Hence,

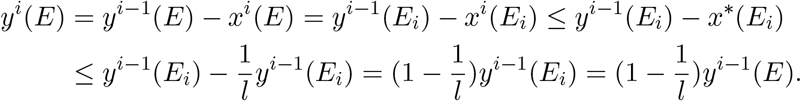

Inductively,

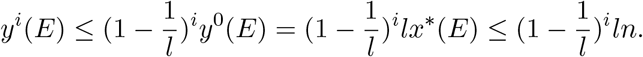

Hence, we have

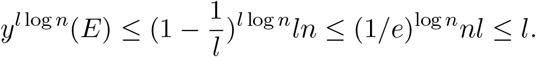

After at most *k*(*k* ≤ *l* log *n*) steps we have 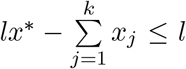. It is easy to see that at most *l* more steps are needed to reach integer *N* such that *lx*\* = *x*^1^ + ⋯ + *x*^N^, where *x_i_* is an incident vector of some feasible matching. Hence, we have

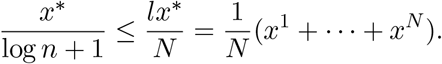

By an averaging argument, there exists an *x^i^* such that 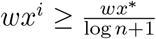.

#### Proof of Theorem 5.

For *u* ∈ *V*_1_ and *v* ∈ *V*_2_, let *D*[*u, v*] denote the weight of a maximum *x*\*-weight clique between [*r*_1_, *u*] and [*v, ℓ_i_*], for any leaf *ℓ_i_* in the subtree rooted at *v*. Then

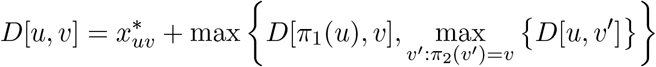

and

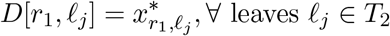

#### Proof of Theorem 6.

For a root-to-leaf path [*r*_1_, *ℓ_j_*] in *T*_1_ we assign weights 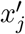 to nodes *v* in *T*_2_ as follows:

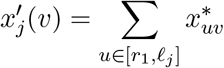

Note that this can be done in 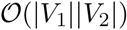. Then, a 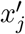-maximum weight independent set of vertices in *T*_2_ gives a maximum semi-independent clique with respect to *P*_1_. Let 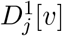 denote the weight of a maximum *x*\*-weight clique between [*r*_1_, *ℓ_j_*] and an independent set in the subtree of *T*_2_ rooted at *v*. Then

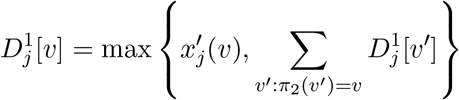

The maximum *x*\*-weight clique 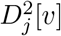 between [*r*_2_, *ℓ_j_*] and an independent set in the subtree of *T*_1_ rooted at *v* is defined analogously in time 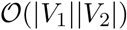. Finally, the maximum weight semi-independent clique constraint can be computed as

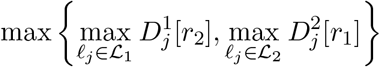

### 6.2 Heuristic approaches implemented in Trajan

A simple *greedy approach* adds the best, i.e., lowest cost edge to a set of already selected edges that is not in conflict with any of them. This strategy can be applied to large scRNA-seq sample comprising hundreds of thousands of cells. Since it completely ignores any optimal fractional solution, it can return very suboptimal solutions.

In contrast, a *randomized rounding* scheme takes into account the global dependence of variables in the LP relaxation: A fractional variable 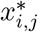 in the optimal solution to the improved LP relaxation is rounded to 1 with probability *x_i,j_*, such that the expected objective function value of the integral solution is the same as of the optimal fractional solution. Potentially introduced conflicts are subsequently resolved by greedily retaining the most valuable (with respect to *w*(*i, j*)) edges. The tighter the LP relaxation the better the integral solution will be.

Another approach is to add the following *non-linear constraints* to the LP to force the variables *x_i,j_* to be either 0 or 1:

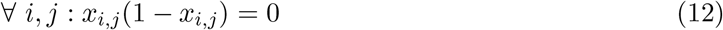

From a high level point of view these integrality constraints can be thought of as a rank-1 approach to the first level of the Lasserre hierarchy [14, 15]. When trying to solve this NP-hard problem using our non-linear solver (Section 6.3), we obtain integral, but suboptimal solutions. We expect this method to return slightly better solution than the randomized rounding scheme, since in contrast to the latter, it enforces integrality at the same time as it solves the relaxed, fractional problem.

### 6.3 Non-linear solver

We use our non-linear solver to solve all continuous optimization problems, unless stated otherwise. It implements the standard augmented Lagrangian approach [12, 17] in order to deal with nonlinear constraints. The augmented Lagrangian approach runs in iterations and maps the constrained optimization problem to a sequence of unconstrained optimization problems that can have bounds on the variables. These unconstrained optimization problems are then solved by the quasi-Newton solver (L-BFGS-B [16, 25]) that can also deal with bounds on the variables. A detailed description on the individual parts of this quasi-Newton solver can be found in [25].

It has been observed before [2] that when the problem is non-convex an augmented Lagrangian approach can be superior to interior point methods with respect to the solution quality. It often returns better local minimal solutions than an interior point solver. Note, that this is the situation when adding the non-linear integrality constraints (12).

### 6.4 Supplemental Figure

**Figure 8:**
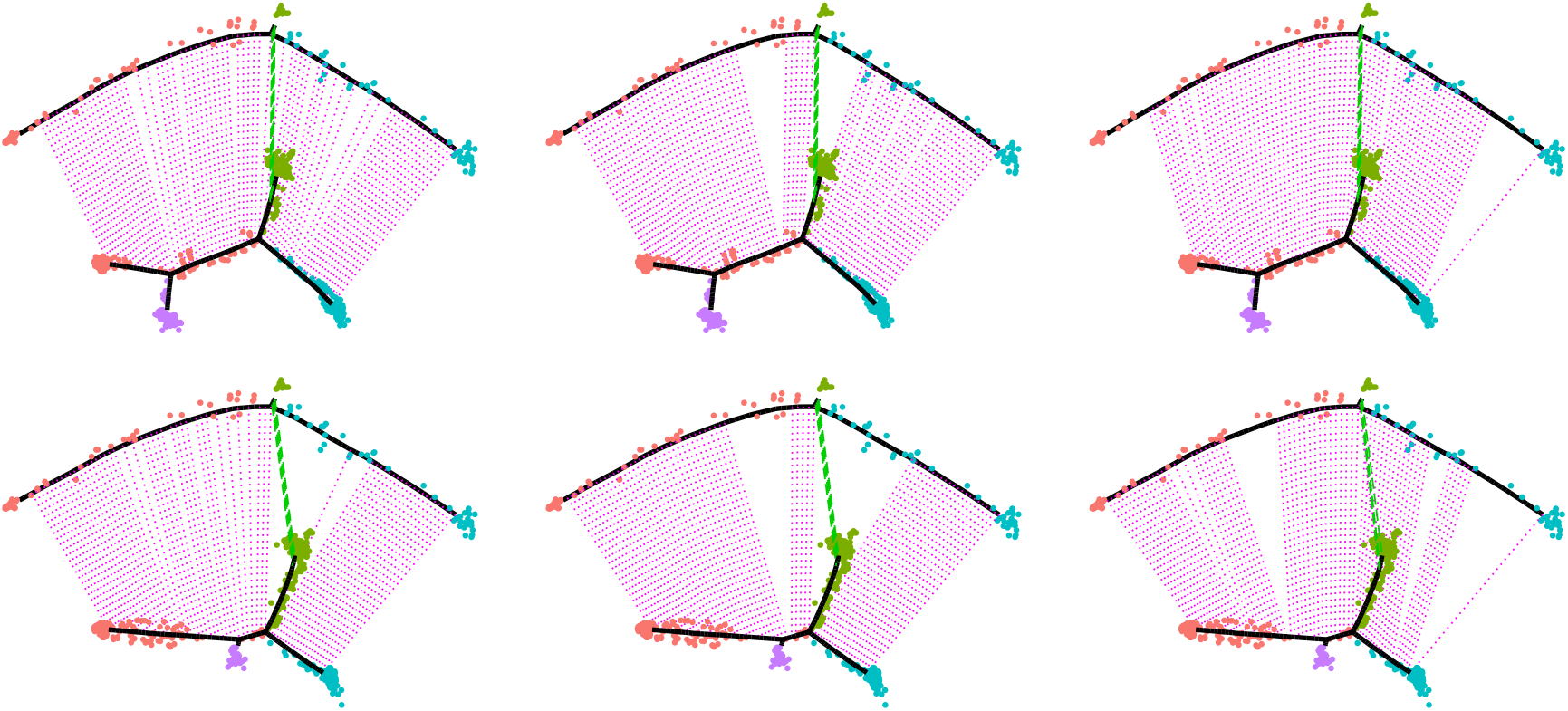
Alignment of myogenic reprogramming and differentiation dynamics using three different penalty schemes, from left to right: avg, max, lb. The bottom row shows results for random subsamples of input cells and ordering genes.

### 6.5 Arboreal DAG matchings

In this section, we generalize arboreal matchings between rooted trees to directed acyclic graphs (DAGs) *G*_1_ = (*V*_1_, *E*_1_) and *G*_2_ = (*V*_2_, *E*_2_). As for trees, we define pairs of edges (*i, j*) and (*k, l*) between *G*_1_ and *G*_2_ to be *compatible* in an arboreal matching if they preserve the ancestry relationship, that is, if it holds that *k* is a descendant of *i* in *T*_1_ if and only if *l* is a descendant of *j* in *T*_2_. Adjusting the definition of set 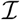 of compatible edges to the descendants relationship given by a DAG rather than a tree, ILP (P) defines the maximum weight arboreal matching between DAGs.

The lifted clique constraints (9) and (11) are analogously valid for DAGs, but their separation needs to be adapted. For crossing clique constraints, the dynamic program can be generalized to DAGs by considering all parents of node *u* (i.e., *π*_1_(*u*) and *π*_2_(*v*′) here stand for sets of nodes) instead of the unique parent in a tree.

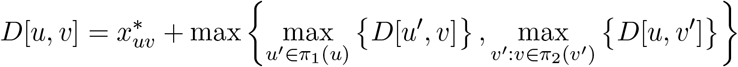

and

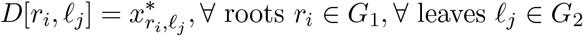

The running time of this DP is 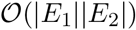 and thus remains polynomial.

The semi-independent clique constraints need to be computed for each root-to-leaf path in *G*_1_ and *G*_2_. For a given root-to-leaf path in *G*_1_ (or *G*_2_), the independent set in *G*_2_ (or *G*_1_) can be represented as a maximum weight antichain which can be reduced to a standard max-flow problem (see [10]), and hence can be computed efficiently. Since the number of paths in a DAG can be exponential, our approach to separate clique constraints is exponential in the worst case. In fact, we show below that we cannot hope for a polynomial time separation algorithm (assuming **P** ≠ **NP**). Next, we formally state the problem of identifying the most violated (maximum) semi-independent clique between two DAGs and show that it is NP-hard.

**Problem**. *Maximum Semi-independent Clique* (MSC)

*Given two DAGs G*_1_ = (*V*_1_, *E*_1_) *and G*_2_ = (*V*_2_, *E*_2_) *with weights α*: *V*_1_ × *V*_2_ → ℝ_+_. *Find a maximum semi-independent clique with respect to the weights α, that is, find a root-to-leaf path P*\* *in G*_1_ *and an independent set S*\* *in G*_2_ *such that the total weight* 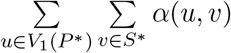 *is maximized*.

Here the weights *α* correspond to the current fractional solution and our goal is to find the most violated semi-independent clique. We prove NP-hardness of MSC via a reduction from the *maximum label cover* problem [4]. Given a bipartite graph *G* = (*A, B, E*), with a partition of *A* and *B* into *k* disjoint sets *A*_1_,…,*A_k_* and *B*_1_,…,*B_k_*, respectively, the maximum label cover problems is to find subsets of vertices *A*′ ⊆ *A* and *B*′ ⊆ *B*, such that, |*A*Ȳ ∩ *A_i_*| ≤ 1 and |*B*′ ∩ *B_i_*| ≤ 1, for *i* = 1,…,*k*, so as to maximize the number of edges

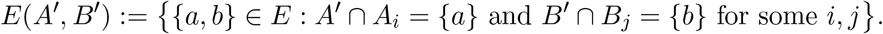

#### Theorem 7.

MSC *is NP-hard*.

*Proof*. Starting from an instance of the maximum label cover problem *G* = (*A, B, E*) and *k* disjoint sets *A*_1_,…, *A_k_* and *B*_1_,…, *B_k_*, we construct a corresponding instance of MSC problem as follows. For every subset *A_i_*, we define a directed graph *D_i_* = (*N_i_, A*(*D_i_*)), where *N_i_* = {*s_i_, s*_*i*+1_} ∪ *A_i_*, *A*(*D_i_*) = {(*s_i_, a*), (*a, s*_*i*+1_)|*a* ∈ *A_i_*|. The whole graph *G*_1_ = (*V*_1_, *E*_1_) is constructed by concatenating the graphs *D_i_* according to the order of their indices. Figure 9 depicts such a construction. Let *P_i_* be a directed path whose vertex set is *B_i_*. The graph *G*_2_ = (*V*_2_, *E*_2_) is obtained by connecting a vertex *s* to one of the endpoints of *P_i_, i* = 1,…, *k* (see Figure 10). The weights *α* between *V*_1_ and *V*_2_ are defined as follows.

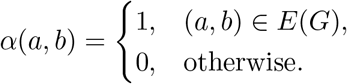

Here, each root-to-leaf path in *G*_1_ (from *s*_1_ to *s*_*k*+1_) corresponds to a selection of exactly one element from set *A_i_, i* = 1,…, *k*, and an independent set in *G*_2_ corresponds to choosing at most one element from set *B_i_, i* = 1,…, *k*. Since only the weight of edges between *a* ∈ *A* and *b* ∈ *B* with (*a, b*) ∈ *E* is nonzero, the maximum semi-independent clique between *G*_1_ and *G*_2_ with respect to the weights *α* is equivalent to the maximum label cover.

**Figure 9:**
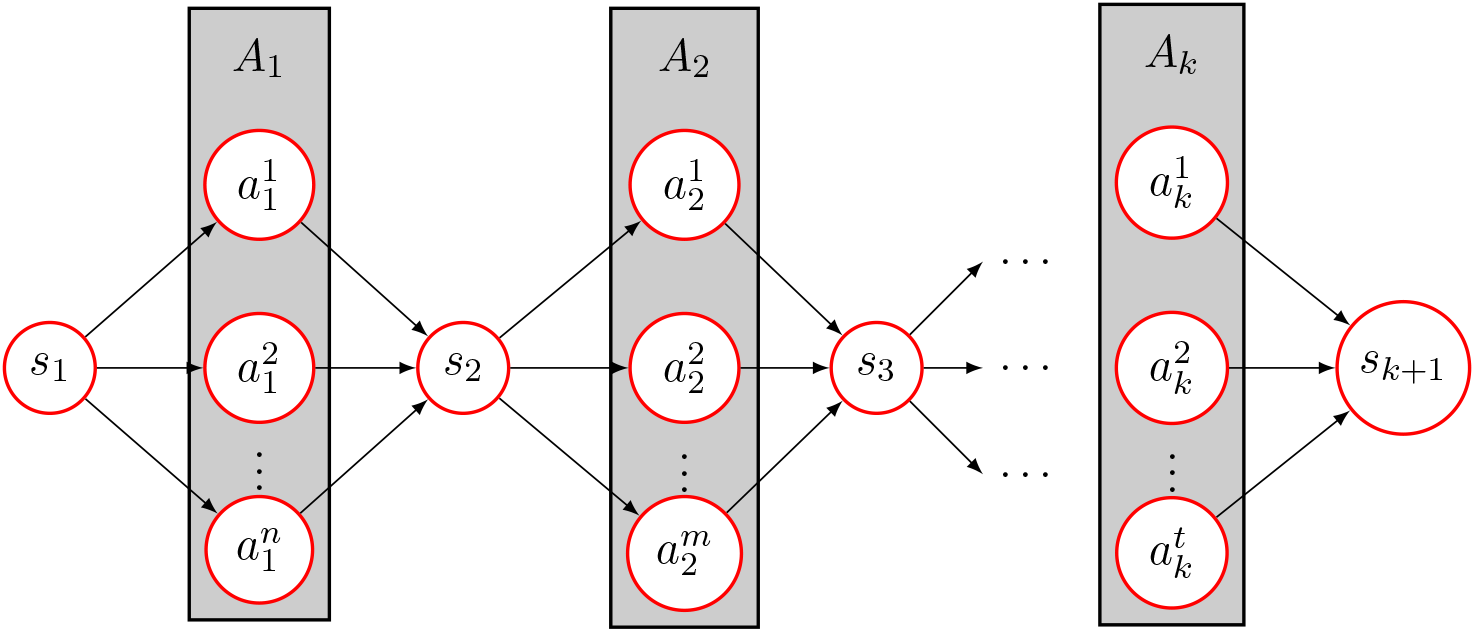
The directed acyclic graph *G*_1_ = (*V*_1_, *E*_1_) associated with an instance *A*_1_,…,*A_k_*. Each gray box contains a set *A_i_*.

**Figure 10:**
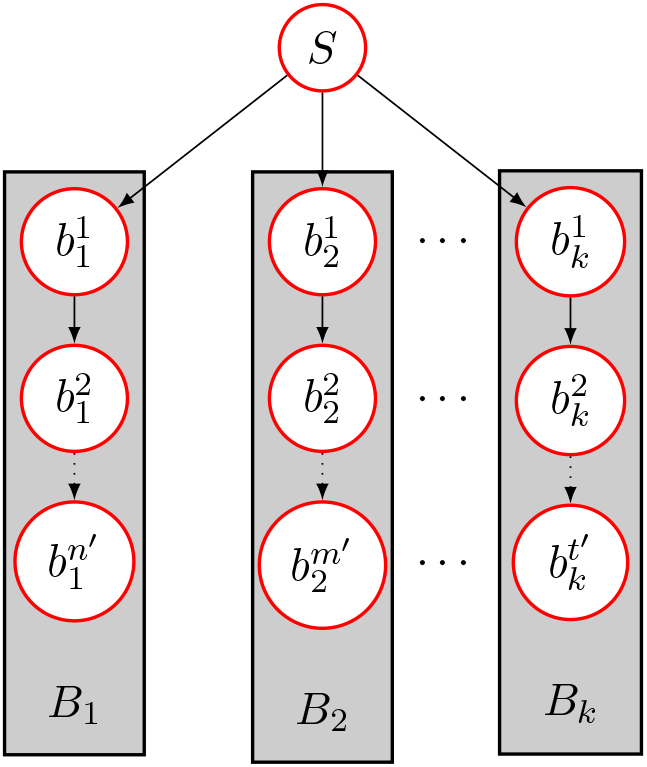
The directed acyclic graph *G*_2_ = (*V*_2_, *E*_2_) associated with an instance *B*_1_,…, *B_k_*. Each gray box contains a set *B_i_*.

